# Elimination of Vesicular Zinc Alters the Behavioural and Neuroanatomical Effects of Social Defeat Stress in Mice

**DOI:** 10.1101/402891

**Authors:** Brendan B. McAllister, David K. Wright, Ryan C. Wortman, Sandy R. Shultz, Richard H. Dyck

## Abstract

Chronic stress can have deleterious effects on mental health, increasing the risk of developing depression or anxiety. But not all individuals are equally affected by stress; some are susceptible while others are more resilient. Understanding the mechanisms that lead to these differing outcomes has been a focus of considerable research. One unexplored mechanism is vesicular zinc – zinc that is released by neurons as a neuromodulator. We examined how chronic stress, induced by repeated social defeat, affects mice that lack vesicular zinc due to genetic deletion of zinc transporter 3 (ZnT3). These mice, unlike wild type mice, did not become socially avoidant of a novel conspecific, suggesting resilience to stress. However, they showed enhanced sensitivity to the potentiating effect of stress on cued fear memory. Thus, the contribution of vesicular zinc to stress susceptibility is not straightforward. Stress also increased anxiety-like behaviour but produced no deficits in a spatial Y-maze test. We found no evidence that microglial activation or hippocampal neurogenesis accounted for the differences in behavioural outcome. Volumetric analysis revealed that ZnT3 KO mice have larger corpus callosum and parietal cortex volumes, and that corpus callosum volume was decreased by stress in ZnT3 KO, but not wild type, mice.

## 1. INTRODUCTION

The physiological and psychological response to stress is often beneficial, helping an organism to meet environmental challenges it might otherwise fail to overcome. Yet the stress response can also be harmful, particularly when the stress is chronic or recurring. Stressful life events increase the likelihood of suffering from mental disorders such as depression (Kendler et al. 1999; Kendler & Gardner, 2016) and anxiety (Kendler et al., 2003). Because of this, the effects of stress on the rodent brain have been widely studied, with the ultimate goal of understanding the mechanisms by which stress contributes to human mental illness.

One way to induce stress in rodents is by exposing them repeatedly to social defeat by a conspecific. A common protocol, developed in the late 1980s (Kudryavtseva et al., 1991), involves subjecting a male mouse to brief, daily episodes of defeat by a series of dominant, aggressive mice. Between defeats, the mouse and its aggressor are housed in close quarters but separated by a partition, allowing sensory contact but preventing fighting or injury. This procedure results in a syndrome of behavioural changes that resembles human depression, including avoidance of other mice (similar to social withdrawal), decreased preference for sucrose (similar to anhedonia), increased anxiety, a sensitized endocrine response to acute stress, and altered circadian rhythms (Berton et al., 2006; Krishnan et al., 2007). Mice that demonstrate these changes are often referred to as susceptible to stress. But a certain proportion are more resilient, exhibiting some of the same changes as susceptible mice but behaving in other respects more similarly to non-defeated controls (Krishnan et al., 2007). This variability in outcome reflects what is seen in humans – not all people who experience stress go on to develop mood disorders (Sheerin et al., 2018) – and provides a model with which to probe the biological factors that predispose an animal toward susceptibility or resilience.

Many such mechanisms have been discovered (Han & Nestler, 2017), but one potential mechanism that has yet to be examined is vesicular zinc. “Vesicular zinc” refers to zinc ions that are sequestered in the synaptic vesicles of neurons (McAllister & Dyck, 2017) – in the forebrain, zinc is found in a subset of glutamatergic neurons (Beaulieu et al., 1992; Sindreu et al., 2003). This zinc is released in an activity-dependent manner and can modulate a plethora of targets, including glutamate receptors (Paoletti et al., 2009; Vergnano et al., 2014; Anderson et al., 2015; Kalappa et al., 2015). Notably, vesicular zinc storage is the responsibility of a protein called zinc transporter 3 (ZnT3; Palmiter et al., 1996; Wenzel et al., 1997), encoded by the *SLC30A3* gene. When this transporter is eliminated, as in the ZnT3 knockout (KO) mouse, vesicular zinc can no longer be detected (Cole et al., 1999), providing a useful tool with which to study the function of vesicular zinc in the brain.

Despite the prevalence of vesicular zinc in the forebrain, mice that lack ZnT3 and vesicular zinc do not show a strong behavioural phenotype (Cole et al., 2001; Thackray et al., 2017). However, mounting evidence indicates that these mice are subtly abnormal. They perform normally when tested using a standard fear conditioning protocol but show deficient learning in a “weaker” training paradigm (Martel et al., 2010). They can perform an object recognition task when the interval between training and testing is short (Wu & Dyck, 2018), but not when it is extended to longer times (Martel et al., 2011). And ZnT3 KO mice can discriminate between textures when the difference is pronounced, but they lack the ability to detect fine textural differences (Wu & Dyck, 2018).

This emerging pattern of behavioural deficits under challenging or complex conditions raises the question of how ZnT3 KO mice respond to the challenge of chronic stress. There is reason to believe that vesicular zinc signaling could modulate stress outcomes. It has not been confirmed whether co-release of zinc occurs in the glutamatergic pathways that modulate the behavioural response to social defeat stress – e.g., ventral hippocampus to nucleus accumbens (NAc), prefrontal cortex (PFC) to NAc, and PFC to amygdala (Kumar et al., 2014; Bagot et al., 2015). But vesicular zinc-containing axon terminals are abundant in these brain regions (Pérez-Clausell & Danscher, 1985; Frederickson et al., 1992), and zinc-containing neurons are known to form reciprocal connections between PFC and amygdala (Christensen & Frederickson, 1998; Cunningham et al., 1997). Given this, we sought to characterize how ZnT3 KO mice respond to repeated social defeat (RSD), to understand whether vesicular zinc influences the outcomes of chronic stress.

## 2. MATERIAL AND METHODS

### 2.1 Animals

All protocols were approved by the Life and Environmental Sciences Animal Care Committee at the University of Calgary and followed the guidelines for the ethical use of animals provided by the Canadian Council on Animal Care. All efforts were made to minimize animal suffering, to reduce the number of animals used, and to utilize alternatives to *in vivo* techniques, if available. Mice were housed in temperature-and humidity-controlled rooms on a 12:12 light/dark cycle (lights on during the day). Food and water were provided *ad libitum*. WT and ZnT3 KO mice, on a mixed C57BL/6×129Sv background, were bred from heterozygous pairs. Offspring were housed with both parents until P21, at which point they were weaned and housed in standard cages (28 × 17.5 × 12 cm with bedding, nesting material, and one enrichment object) in groups of 2-5 same-sex littermates. CD-1 mice used for the RSD procedure were retired breeders, 4-12 months old, from the University of Calgary Transgenic Services Facility or Charles River.

### 2.2 Experimental design

For a diagram depicting the experimental design, see Figure 1. At 8-10 weeks of age, WT and ZnT3 KO mice were assigned either to the stress or control condition. Stress consisted of 10 days of RSD (day 1 to day 10), followed by isolated housing for the remainder of the experiment. The control mice remained in standard group-housing throughout the experiment and were handled daily from day 1 to day 10.

**Figure 1.**
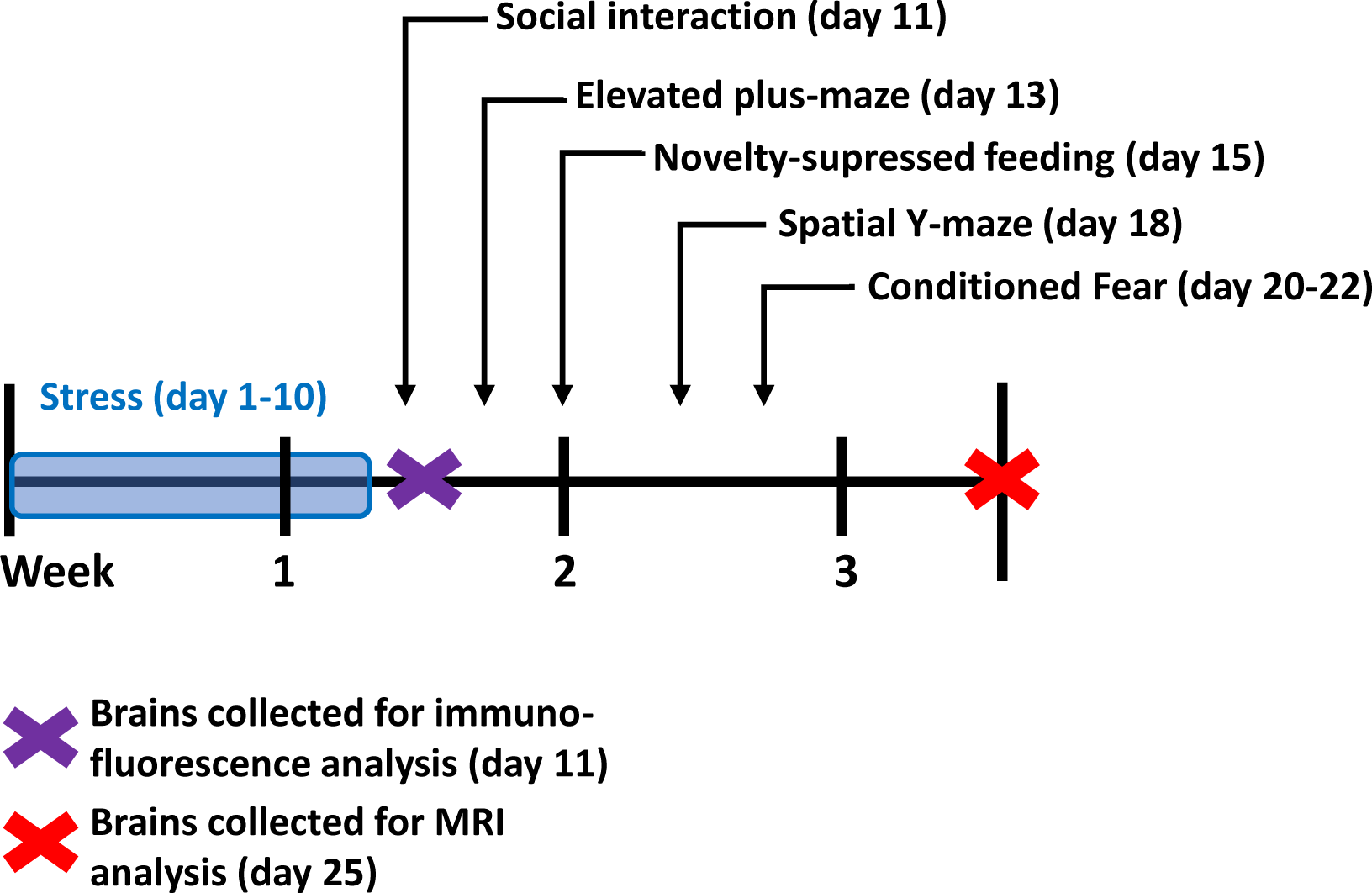
Timeline depicting the experimental design. WT and ZnT3 KO mice were subjected to 10 days of stress, consisting of daily episodes of social defeat, followed by isolated housing for the remainder of the experiment. Control WT and ZnT3 KO mice remained in group housing with their same-sex littermates throughout the experiment.

One day post-RSD (day 11), all mice were subjected to social interaction testing (WT-control: *n* = 20; WT-stress: *n* = 22; KO-control: *n* = 22; KO-stress: *n* = 22). Following this, one cohort of mice was killed, and their brains were extracted for immunofluorescence analysis (*n* = 6 per group). The remaining mice were subjected to further behavioural testing (WT-control: *n* = 14; WT-stress: *n* = 16; KO-control: *n* = 16; KO-stress: *n* = 16). On day 25, brains were extracted from a subset of these mice, to be used for magnetic resonance imaging (MRI) volumetric analysis (WT-control: *n* = 8; WT-stress: *n* = 9; KO-control: *n* = 9; KO-stress: *n* = 10).

### 2.3 Repeated social defeat

The RSD procedure was adapted from Golden et al. (2011). Mice were subjected to daily episodes of defeat for 10 days. For each defeat, the mouse was transferred to a novel CD-1 mouse’s cage for a period of 5 min. During this time, the CD-1 resident would reliably attack the smaller intruder. After three attacks (with an attack defined as an uninterrupted episode of physical interaction, almost always resulting in vocalizations from the intruder), the intruder was placed in a mesh enclosure (8.5 cm Ø) for the remainder of the 5 min, allowing the mice to interact in close proximity but restricting further fighting or injury. Following this, the intruder was housed with the CD-1 resident, but with the two mice separated by a perforated acrylic partition that divided the large cage (24 × 45.5 × 15 cm) lengthwise into two compartments, allowing for visual, auditory, and olfactory contact, but restricting physical interaction. Prior to each defeat, the intruder mice were rotated between cages, in order to prevent them from habituating to a particular CD-1 resident. After the final defeat, the mice were singly-housed in standard cages. CD-1 mice were prescreened for aggressiveness, as described by Golden et al. (2011).

### 2.4 Behavioural assessment

Testing was conducted during the light phase, and the mice were allowed to habituate to the behavioural testing room for 30 min prior to the start of each test. The behavioural tests were conducted in the order presented below.

#### 2.4.1 Social interaction

The procedure for the social interaction test was adapted from Golden et al. (2011). The test was conducted under dim red light. The apparatus for the test was an open field (40 × 40 cm) constructed of white corrugated plastic. The test consisted of three 150 s phases, each separated by 60 s. For the first phase, a mesh enclosure (10 cm Ø) was placed against a wall of the field; the mouse was then placed along the center of the opposing wall and allowed to explore freely. The second phase was the same, but with a novel, age-matched mouse of the same strain (novel conspecific) placed inside the enclosure. For the third phase, the conspecific was replaced by a novel, aggressive CD-1 mouse. Between testing each mouse, the enclosures and the field were cleaned with Virkon; the field was also cleaned of urine and feces between each phase. The test was recorded using a digital video camera with night-vision capability (Sony HDR-SR8), and scoring was automated (ANY-maze, version 4.73). The following parameters were scored: “interaction time” (i.e., time in the interaction zone, defined as a 26 × 16 cm rectangle around the enclosure); “corner time” (i.e., time in either of the two corners of the field opposing the enclosure, each encompassing a 9 × 9 cm area); total distance traveled; and “immobility time” (minimum period of 2 s, with detection sensitivity set at 90%). Social interaction ratios were calculated by dividing interaction time in the third phase by interaction time in the first phase.

#### 2.4.2 Elevated plus-maze

The elevated plus-maze (EPM) test was conducted as previously described (McAllister et al., 2015), with minor modifications. The apparatus was a plus-shaped structure, with two opposing open arms, two closed arms, and a center where the arms intersected. The maze was illuminated by dim light (3 lux). Mice were tested for 5 min. Activity was video recorded and scoring was automated using ANY-maze. The following parameters were scored: open arm time, center time, and total distance traveled. The amount of time spent on the open arms was used as an indicator of anxiety, with less time assumed to reflect greater anxiety.

#### 2.4.3 Novelty-suppressed feeding

The protocol for the novelty-suppressed feeding (NSF) test was adapted from Samuels and Hen (2011). Mice were food deprived for 16 h prior to the test. The test was conducted in an open field (40 × 40 cm) under bright lighting (800 lux). The floor of the field was covered with wood-chip bedding. A food pellet was fixed to a small platform in the center of the field, preventing the mouse from moving the pellet. The latency to begin feeding was recorded, up to a maximum time of 10 min. The mouse was then returned to its home-cage (with its cage-mates temporarily removed) and transported immediately to an adjacent, dimly lit (3 lux) room. A pre-weighed food pellet was placed in the hopper, and the latency to begin feeding was recorded; a maximum score of 180 s was given if the latency exceeded that length. Once the mouse began feeding it was allowed 5 min to eat, after which the pellet was removed and weighed, to calculate food consumption. Body weight was also recorded both prior to food deprivation and after the test. Longer latencies to feed in the novel field were assumed to reflect greater anxiety.

#### 2.4.4 Spatial Y-maze

The Y-maze protocol was adapted from Conrad et al. (2003). The apparatus was a three-armed wooden structure, painted black. Each arm measured 10.5 by 48 cm, with 15 cm high walls. The structure was elevated off the ground, and there were numerous visual cues surrounding the maze to be used for spatial orientation. The floor was covered in bedding, which was mixed between trials to prevent odours being used as non-spatial cues. For the training phase of the test, mice were placed at the end of the “starting arm” and allowed to explore for 15 min, with one arm blocked off by a partition. The mice were returned to the maze 3 h later for a 5 min test phase, in which the partition was removed, providing a novel arm that had not previously been explored. The test phase was video recorded for scoring. Because mice have a natural propensity to explore novel areas, it was assumed that mice with intact spatial memory would make a greater percentage of their total entries into the novel arm in comparison to the “other arm” (i.e., the third arm that was not the starting arm or novel arm). The total number of arm entries was used as an indicator of locomotor activity.

#### 2.4.5 Conditioned fear

A 3-day conditioned fear test was conducted to assess cued and contextual fear memory. The former involves learning an association between an initially neutral stimulus (e.g., a tone) and a noxious stimulus (e.g., an electric footshock), whereas the latter involves forming an association between a noxious stimulus and the context in which it is encountered. The test was performed using a conditioning box (Hamilton-Kinder LM1000-B). Between mice, the inside of the box was wiped with a disinfectant solution. On day 1, after a 2 min period of acclimatization to the box, a 20 s tone was presented. A single footshock (2 s, 0.3 mA) was administered coinciding with the final 2 s of the tone. After an additional 30 s, the mouse was removed from the box. On day 2, the apparatus was altered to provide a novel context (black walls were replaced with white walls, a solid plastic insert was placed over the metal grid floor, coconut scent was added to the chamber, and Virkon was used in place of 70% ethanol as a disinfectant). Mice were allowed to explore the box for 1 min, after which the tone was presented for 2 min to assess the fear response to the auditory cue (i.e., cued fear memory). On day 3, the apparatus was reverted to its original conditions, and the mice were tested for 3 min to assess their fear response to the context (i.e., contextual fear memory). The activity of the mice was video-recorded for analysis of freezing (defined as total immobility, excluding minor movements associated with breathing), which was automated using ANY-maze (minimum freezing duration = 500 ms).

### 2.5 Anatomical analyses

#### 2.5.1 Immunofluorescence labeling

Mice were deeply anaesthetized with an overdose of sodium pentobarbital, and transcardially perfused with phosphate buffered saline (PBS) followed by 4% paraformaldehyde (PFA) in PBS. Brains were extracted and post-fixed overnight in 4% PFA in PBS at 4 °C. The spleen and adrenal glands were also extracted and weighed. After post-fixing, the brains were transferred to a sucrose solution (30% sucrose, 0.02% sodium azide in PBS) and stored at 4 °C. Brains were cut coronally into six series of 40 µm sections using a sliding microtome (American Optical, Model #860). One series was labeled for the cellular proliferation marker Ki67 (1:2000, Leica NCL-Ki67p). A second series was labeled for the microglia marker Iba1 (1:1000, Wako #019-19741). A detailed protocol is provided in the supplementary methods. Sections were mounted on gelatin-coated slides, coverslipped with fluorescence mounting medium, and stored at 4 °C.

#### 2.5.2 Hippocampal cell counting

Ki67^+^ cells were counted in each section using an epi-fluorescence microscope (Zeiss Axioskop 2) with a 63×/1.40 objective. Cells were counted in the granule cell layer and the subgranular zone (defined as three cell-widths from the hilar edge of the granule cell layer) of the dentate gyrus. The counts were multiplied by six to estimate the total number of Ki67^+^ cells.

#### 2.5.3 Microglial analysis

Microglia were assessed in the PFC (prelimbic region), basolateral amygdala (BLA), dorsal hippocampus (dHPC; dentate gyrus region), and ventral hippocampus (vHPC; CA3 region). Images were captured bilaterally from three sections, resulting in a total of six images per region of interest (ROI).

Changes in microglial morphology, such as increased soma size and altered process length or number, occur when microglia become “activated.” A thresholding method was used to provide a gross assessment of such changes. This method involves binarizing an image into areas of positive and negative labeling (Beynon & Walker, 2012). Images for analysis were generated by capturing z-stacks throughout the depth of the tissue section, using a confocal microscope (Nikon C1si) with a 20× objective. The “volume render” function of EZ-C1 software (Nikon) was used to collapse the stack into a single image. The images were processed with the “subtract background” and “sharpen” functions of ImageJ (http://rsb.info.nih.gov/ij/), and the “threshold” function was used to binarize the images. The optimal threshold level was determined across several sections from different brains, and then applied uniformly to all the images. The percentage of Iba1^+^ area was then measured.

Microglial density was also quantified. Images were captured using a microscope (Zeiss Axioskop 2) with a 10×/0.30 objective. The number of microglia within each ROI was counted using the “multi-point” tool in ImageJ, then divided by the area of the ROI.

#### 2.5.4 MRI acquisition and analysis

Mice were perfused as described above. Brains were stored in 4% PFA in PBS at 4 °C. MRI acquisition was conducted as previously described (Wright et al., 2017, 2018). Brains were washed overnight in PBS and embedded in 2-3% agar for ex-vivo MRI using a 4.7 T Bruker MRI (Bruker, Ettlingen, Germany Biospin, USA). A 3D multiple gradient echo sequence was acquired using a cryogenically-cooled RF coil and the following imaging parameters: repetition time = 110 ms; echo times = 4, 8, 12…80 ms; matrix = 176 × 128 × 70; field of view = 17.6 × 12.8 × 7 mm^3^; and voxel size = 0.1 mm^3^. Echoes were averaged offline for ROI delineation.

For analysis, six *a priori* ROIs – prefrontal cortex, hippocampus, corpus callosum (CC), parietal cortex, lateral ventricles (LV), amygdala – were traced per hemisphere using ITK-SNAP (www.itksnap.org) as previously described (Shultz et al., 2013; Wright et al., 2017, 2018). “Prefrontal cortex” was defined as all cortex in the 12 slices anterior to the forceps minor. “Hippocampus” started at the anterior tip of the CA3 field and continued for 15 slices (encompassing most of dorsal, but not ventral, hippocampus). Analysis of the remaining structures was also limited to these 15 slices. “Parietal cortex” was defined as all cortex dorsal and lateral to the CC, with the rhinal fissure serving as the ventral boundary. “Amygdala” was defined as everything ventral to the rhinal fissure and lateral to the external capsule and striatum (thus including structures surrounding the amygdala such as the entorhinal and piriform cortex). ROI volumes were determined using Fslutils, a component of FMRIB’s Software Library (FSL, www.fmrib.ox.ac.uk/fsl). MRI analysis was conducted by a researcher blind to the experimental conditions.

### 2.6 Statistical analysis

Statistical analyses were conducted using IBM SPSS Statistics (version 21). Unless otherwise stated, comparisons were conducted by two-way analysis of variance (ANOVA) with genotype (WT vs. ZnT3 KO) and stress (control vs. stress) as factors. Significant interactions were followed-up using Bonferroni-corrected simple-effects tests. All ANOVA results are reported in Supplemental Table 1. Means are presented ± standard deviation.

**Table 1.** Additional behavioural measures. Statistics are reported as mean ± standard deviation. *Main effect of stress, *p* < .05. †Stress × genotype interaction, *p* < .05.

|  | Wild type control | Wild type stress | ZnT3 KO control | ZnT3 KO defeated |
| --- | --- | --- | --- | --- |
| **Social interaction test** | ($n = 20$) | ($n = 22$) | ($n = 22$) | ($n = 22$) |
| Distance (m) – empty cage | 8.9 ± 1.7 | 8.5 ± 1.9 | 8.7 ± 1.8 | 8.7 ± 2.6 |
| Interaction time (s) – empty cage* | 61.3 ± 13.2 | 53.9 ± 18.7 | 59.7 ± 21.1 | 49.4 ± 19.2 |
| Corner time (s) – empty cage | 25.4 ± 7.9 | 27.0 ± 13.0 | 25.3 ± 13.0 | 32.3 ± 23.4 |
| Immobility time (s) – conspecific* | 14.1 ± 11.4 | 39.8 ± 17.4 | 18.5 ± 16.3 | 33.9 ± 15.0 |
| Immobility time (s) – CD-1* | 17.9 ± 8.82 | 42.2 ± 28.2 | 23.9 ± 25.2 | 50.6 ± 36.0 |
| **Elevated plus-maze** | ($n = 14$) | ($n = 16$) | ($n = 16$) | ($n = 16$) |
| Distance traveled (m) | 5.6 ± 1.5 | 5.8 ± 2.0 | 6.2 ± 1.7 | 5.2 ± 1.1 |
| **Novelty-suppressed feeding** | ($n = 14$) | ($n = 16$) | ($n = 16$) | ($n = 16$) |
| Latency to feed (s) – home cage | 31.4 ± 32.9 | 25.9 ± 27.6 | 38.6 ± 39.4 | 31.3 ± 33.1 |
| Food consumption (g) | 1.23 ± 0.09 | 0.09 ± 0.06 | 0.08 ± 0.06 | 0.07 ± 0.06 |
| **Y-maze spatial memory** | ($n = 14$) | ($n = 16$) | ($n = 16$) | ($n = 16$) |
| Novel arm entries (% of total entries) | 40.3 ± 6.2 | 42.0 ± 4.4 | 40.2 ± 6.3 | 40.5 ± 5.0 |
| Novel arm time (s) | 100.8 ± 31.3 | 94.9 ± 32.6 | 101.9 ± 28.4 | 92.6 ± 33.5 |
| Total arm entries† | 14.4 ± 6.8 | 16.8 ± 5.4 | 17.9 ± 5.1 | 14.5 ± 4.8 |

## 3. RESULTS

### 3.1 Behavioural assessment

#### 3.1.1 Social interaction

Time spent in the interaction zone and in the corners of the field was examined for the three phases of the test (Figure 2A; Table 1). In the first phase (empty cage), stress decreased interaction zone time by 15% [*F*(1,82) = 4.92, *p* = .029], with no difference between genotypes. For corner time, there was no effect of stress or genotype.

**Figure 2.**
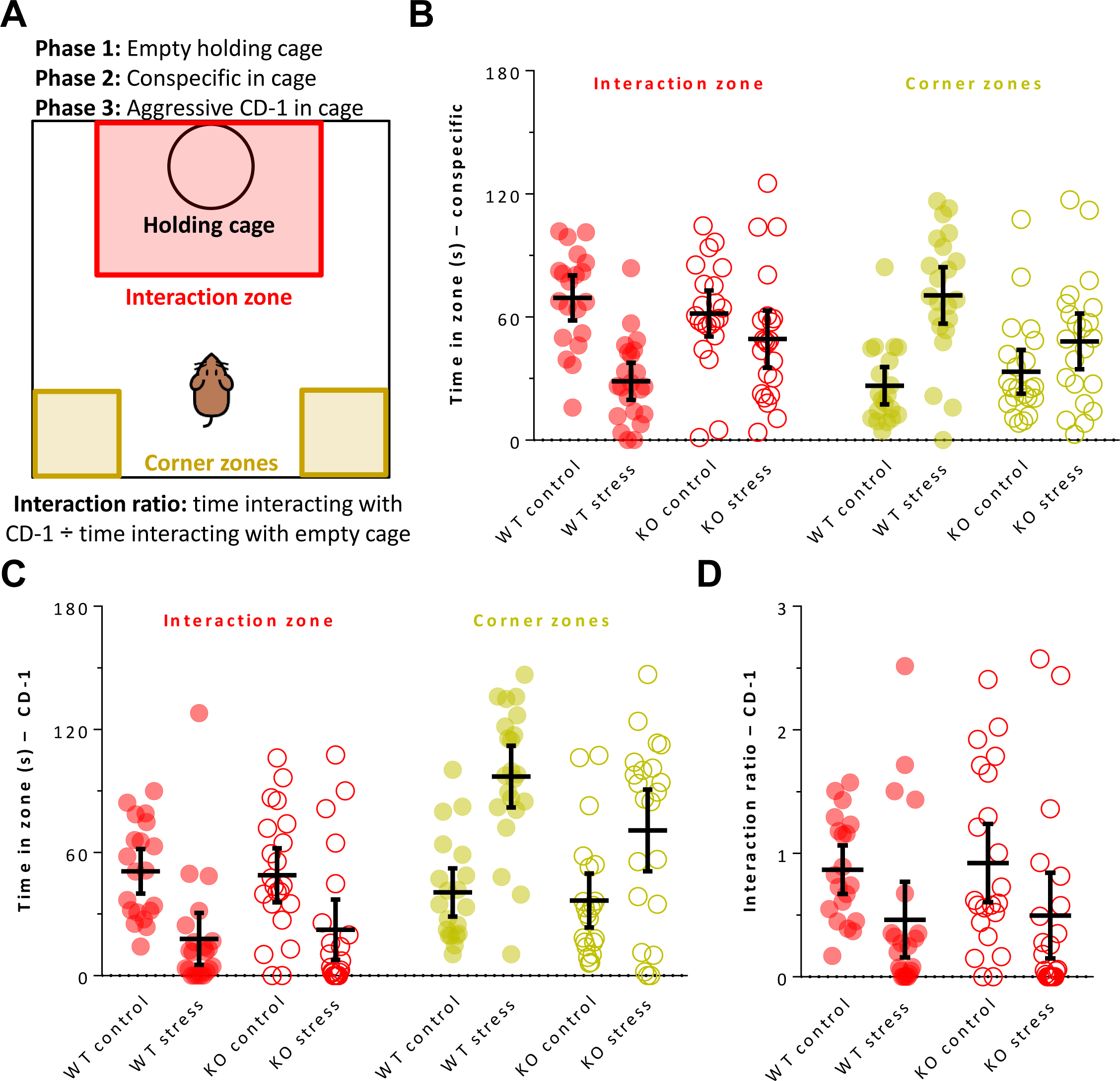
Social interaction behaviour of WT and ZnT3 KO mice following repeated social defeat stress. **A.** Diagram of the social interaction apparatus and explanation of the three phases of the test, each lasting 2.5 min. **B.** Time spent in the interaction zone and corner zones with a novel conspecific in the holding cage (phase 2). Stressed WT mice spent less time in the interaction zone, and more time in the corners, than control mice, whereas stressed ZnT3 KO mice did not differ from controls. **C.** Regardless of genotype, stress decreased time in the interaction zone and increased time in the corner zones when a novel aggressive CD-1 mouse was in the holding cage (phase 3). **D.** Stress decreased the social interaction ratios of the mice. Error bars represent 95% CIs.

In the second phase (novel conspecific; Figure 2B), stress had differing effects on interaction time depending on the genotype of the mice [stress × genotype interaction: *F*(1,82) = 6.53, *p* = .012]. Stress decreased interaction time by 59% in the WT mice (*p* < .001, Bonferroni-corrected], but did not significantly affect the ZnT3 KO mice (*p* = .219). Similarly, there was a significant interaction for corner time [*F*(1,82) = 6.36, *p* = .014]. Stress more than doubled the time spent in the corners by the WT mice (*p* < .001, Bonferroni-corrected), but did not significantly affect the ZnT3 KO mice (*p* = .071). Together, these results indicate that stress caused WT mice to become socially avoidant of a novel conspecific, while ZnT3 KO mice were unaffected.

For the third phase (CD-1 aggressor; Figure 2C), stress decreased interaction time by 60% [*F*(1,82) = 22.53, *p* < .001], and increased corner time by 118% [*F*(1,82) = 37.18, *p* < .001]. Interaction time with the CD-1 mouse did not differ between genotypes, but there was a difference in corner time [*F*(1,82) = 4.12, *p* = .046], with the WT mice spending more time in the corners than the ZnT3 KO mice. To summarize, while only WT mice were avoidant of a novel conspecific following stress, stress caused both WT and ZnT3 KO mice to avoid a novel CD-1 mouse.

There were no differences in total distance traveled in the field during phase 1 of the test (Table 1), indicating that baseline differences in locomotion did not affect the results. During phase 2, stress more than doubled the amount of time spent immobile [*F*(1,82) = 38.72, *p* < .001; Table 1], with no difference between genotypes. The same was true for phase 3 [*F*(1,82) = 19.53, *p* < .001; Table 1].

The social interaction ratio with a CD-1 mouse is the standard measure used to define susceptibility to stress in the RSD model, so we also compared the groups on this measure. One mouse was excluded from this analysis because it spent no time in the interaction zone during phase 1, which prevented us from calculating a ratio. Stress decreased the interaction ratio by 47% [*F*(1,81) = 8.32, *p* = .005; Figure 2D], with no significant difference between genotypes.

#### 3.1.2 Elevated plus-maze

Stress increased anxiety-like behaviour, as indicated by the stressed mice spending less time – indeed, almost no time at all – on the open arms [*F*(1,58) = 8.74, *p* = .004; Figure 3A]. There was no difference between genotypes. Stress also decreased the amount of time spent in the center of the maze [*F*(1,58) = 10.99, *p* = .002; Figure 3B]. The ZnT3 KO mice tended to spend less time in the center of the maze than did the WT mice, though this effect was not significant [*F*(1,58) = 3.21, *p* = .078]. There was no effect of stress or genotype on total distance traveled (Table 1), indicating that differences in locomotor activity did not influence the results of this test.

**Figure 3.**
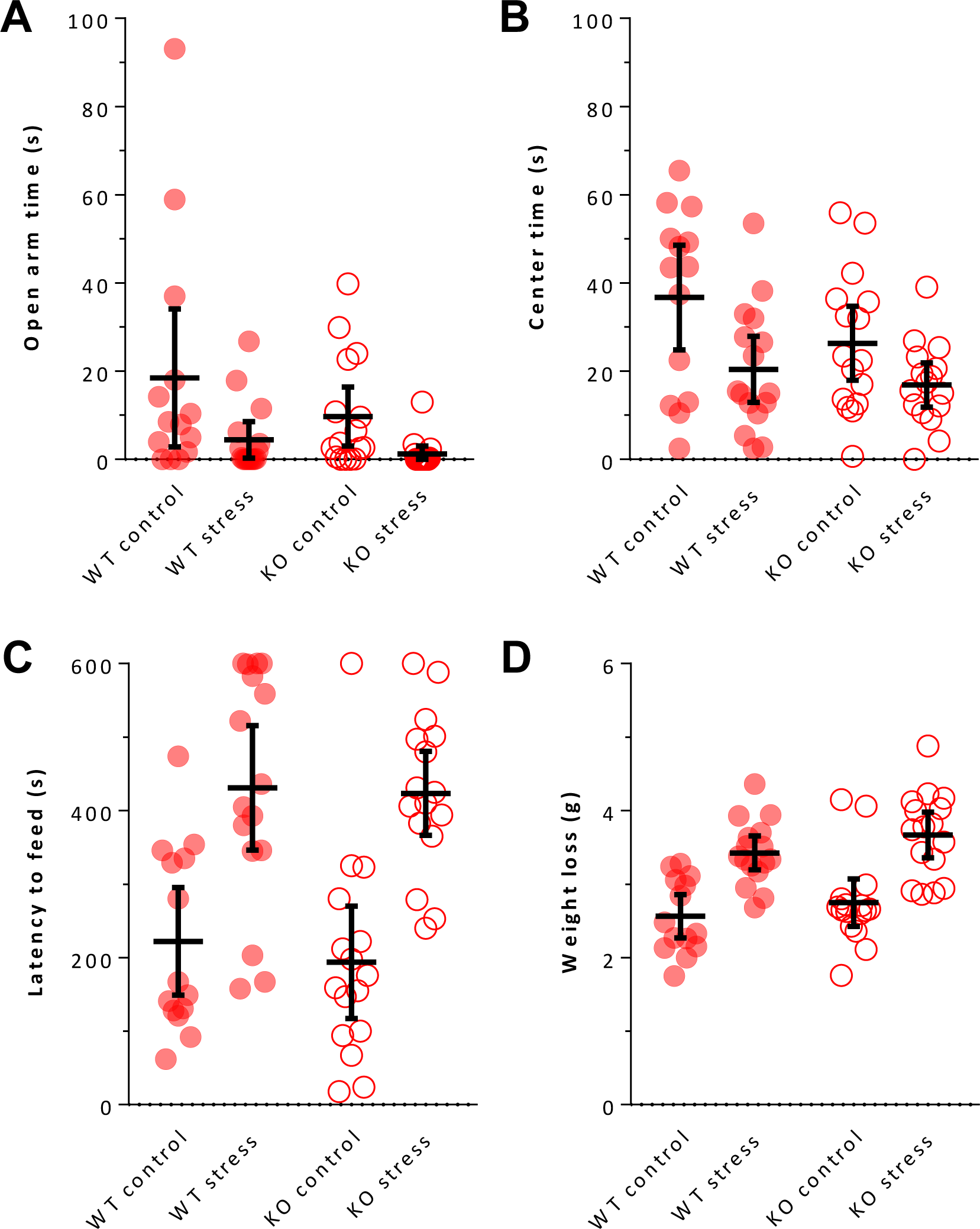
Anxiety-like behaviour of WT and ZnT3 KO mice following repeated social defeat stress. **A.** Stress decreased the time spent on the open arms during a 5 min test in an elevated plus-maze, indicating increased anxiety. **B.** Stress also decreased time spent in the center area of the elevated plus-maze. **C.** Regardless of genotype, stress increased the latency to begin feeding when food-deprived mice were placed in a novel environment, again indicating increased anxiety. The maximum allowed time was 10 min. **D.** Relative to controls, stressed mice lost more weight over the 16 h food deprivation period prior to the novelty-supressed feeding test. Error bars represent 95% CIs.

#### 3.1.3 Novelty-suppressed feeding

Providing further evidence of anxiety-like behaviour, stress increased the latency to begin feeding in the novel open field [*F*(1,58) = 40.43, *p* < .001; Figure 3C], with the stressed mice taking more than twice as long to begin feeding than controls. There was no difference between genotypes. In the home cage, there was no significant difference in the latency to feed between the stressed and control mice [*F*(1,58) = 0.56, *p* = .459; Table 1], suggesting that the difference in the novel field was due to increased sensitivity to the anxiogenic environment, and not due to general changes in feeding behaviour. As further support of this, there was no effect of stress on the amount of food consumed in the home cage [*F*(1,58) = 1.06, *p* = .307; Table 1]. The WT mice did tend to consume more than the ZnT3 KO mice, though this difference was not significant [*F*(1,58) = 3.68, *p* = .060].

Interestingly, there was a significant effect of stress on the change in body weight over the 16 h food restriction period [*F*(1,58) = 42.65, *p* < .001], with the stressed mice losing 33% more weight than the control mice. There was no difference between genotypes.

#### 3.1.4 Spatial Y-maze

Stress did not alter the percentage of total arm entries made into the novel arm or the amount of time spent in the novel arm, nor was there an effect of genotype on either measure (Table 1). There was a significant interaction between stress and genotype on the total number of arm entries [*F*(1,58) = 4.18, *p* = .045; Table 1], with stress tending to increase arm entries in the WT mice (*p* = .244) and decrease arm entries in the ZnT3 KO mice (*p* = .090), but neither effect was significant.

Wilcoxon signed-rank tests were used to compare the percentage of entries into the novel arm to the percentage of entries into the “other arm” within each group. All four groups made a significantly greater percentage of arm entries into the novel arm than the “other arm”, indicating that all groups could remember which arm had been inaccessible during the training phase 3 hours prior (WT-control: 40.3 ± 6.2 vs. 31.1 ± 5.2, *Z* = 2.50, *p* = .013; WT-stress: 42.0 ± 4.4 vs. 28.7 ± 5.9, *Z* = 3.30, *p* = .001; KO-control: 40.2 ± 6.3 vs. 31.7 ± 4.5, *Z* = 2.66, *p* = .008; KO-stress: 40.5 ± 5.0 vs. 32.9 ± 5.8, *Z* = 2.51, *p* = .012).

#### 3.1.5. Conditioned fear

There were no significant effects of genotype or stress on the percentage of time spent freezing before, during, or after the presentation of the tone/shock on day 1 of the test (*p* > .15 for all comparisons; data not shown), indicating that there were no baseline differences in fear or freezing behaviour, and no differences in the initial response to the tone/shock.

Cued fear memory was assessed on day 2 in a novel context. There was no effect of stress or genotype on freezing time before cue presentation (Figure 4A), indicating no baseline differences in the fear response to the novel context. Stress had differing effects on cued fear memory depending on the genotype of the mice [stress × genotype interaction: *F*(1,58) = 5.89, *p* = .018; Figure 4A]. In the ZnT3 KO mice, stress enhanced fear memory, as indicated by increased freezing time during the cue presentation (*p* = .004, Bonferroni-corrected). In the WT mice, stress had no significant effect (*p* = .752). However, it also appeared that control ZnT3 KO mice showed weaker fear memory than control WT mice, as was previously reported by Martel et al. (2010). Therefore, we also directly compared the control groups, and confirmed that ZnT3 KO mice froze less than WT mice during cue presentation [t-test: *t*(28) = 2.16, *p* = .040], indicating weaker cued fear memory.

**Figure 4.**
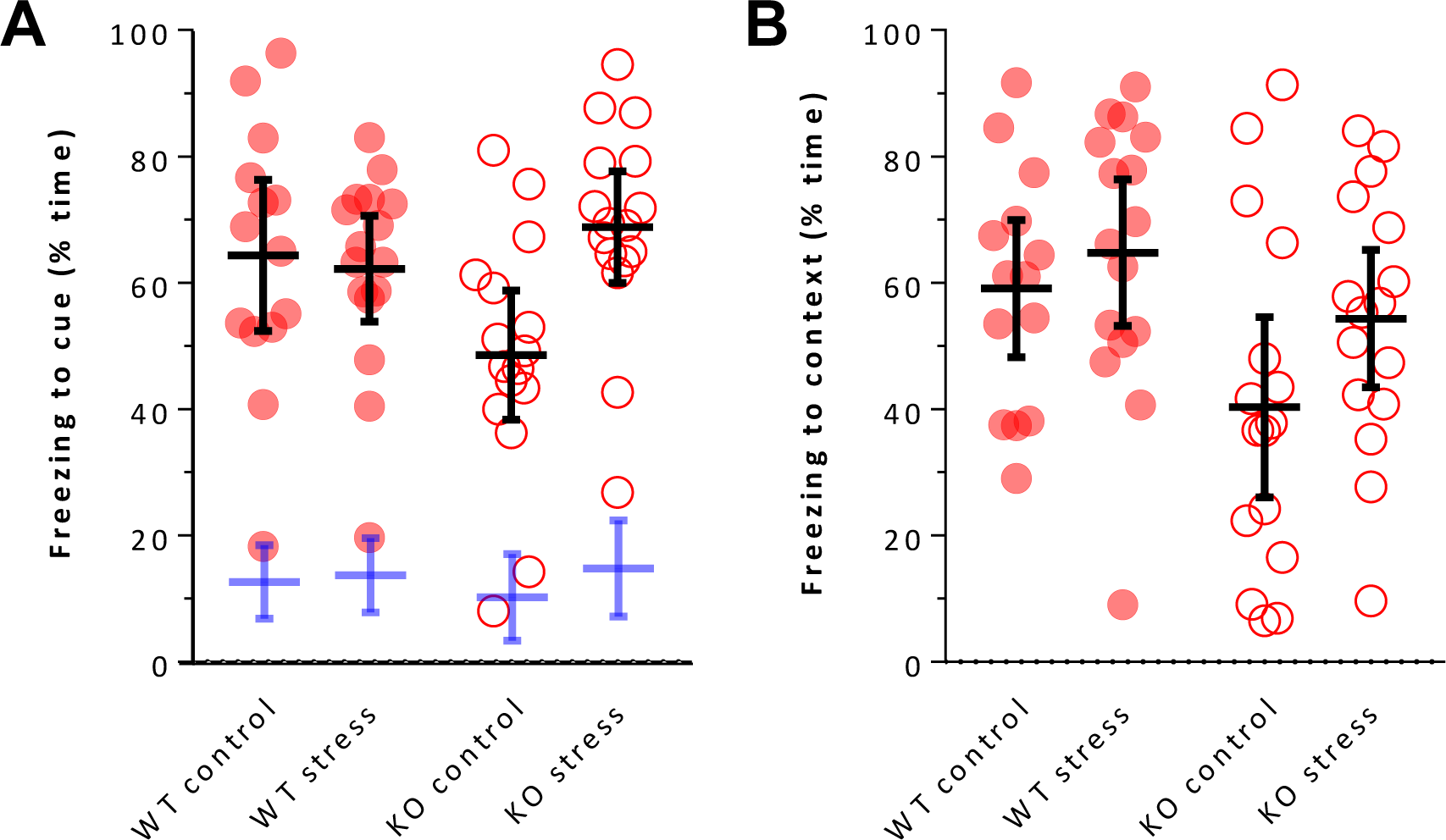
Conditioned fear memory in WT and ZnT3 KO mice following repeated social defeat stress. Training was conducted on day 1, and consisted of a single tone-shock pairing. **A.** On day 2, the freezing response to the tone presented in a novel context (i.e., cued fear memory) was assessed. Cued fear memory was enhanced by stress in ZnT3 KO mice, but was unaffected by stress in WT mice. The blue bars indicate the percentage of time spent freezing in the novel context prior to cue presentation, showing that there were no baseline differences. **B.** On day 3, contextual fear memory was assessed by measuring the time spent freezing when mice were re-exposed to the context in which they were previously shocked. ZnT3 KO mice exhibited less freezing, indicating worse fear memory, than WT mice, regardless of stress. Error bars represent 95% CIs.

Contextual fear memory was assessed on day 3. There was no significant effect of stress on contextual fear memory [*F*(1,58) = 3.02, *p* = .087; Figure 4B], though stress did tend to increase freezing. There was, however, a significant difference between the genotypes [*F*(1,58) = 6.65, *p* = .012], with the ZnT3 KO mice showing weaker contextual fear memory, freezing less than the WT mice.

### 3.2 Body and organ weights

To assess whether stress affected body weight, we calculated weight change from 1 day pre-RSD to 1 day-post RSD. Five mice (2 WT-stress, 2 KO-control, 1 KO-stress) were excluded from this analysis because body weight data were missing from one of the two time points. Neither stress nor genotype influenced the change in body weight. On average, mice gained a small amount of weight over the time period (WT-control: 0.4 ± 1.0 g; WT-stress: 0.2 ± 1.5 g; KO-control: 0.4 ± 1.0 g; KO-stress: 0.3 ± 1.0 g).

Spleen and adrenal weights were also examined in the mice killed 1 day following stress (Table 2). Stress significantly increased spleen weight [*F*(1,20) = 5.76, *p* = .026], but there was no difference between genotypes. When spleen weights were analyzed as a percentage of total body weight, the same pattern of findings was observed (data not shown). Stress also significantly increased adrenal gland weight [*F*(1,20) = 15.86, *p* < .001], with no difference between genotypes. However, when adrenal weight was analyzed as a percentage of total body weight, the effect of stress was found to differ between genotypes [stress × genotype interaction: *F*(1,20) = 7.87, *p* = .011]. Stress increased adrenal weight in the ZnT3 KO mice (*p* < .001, Bonferroni-corrected), but not in the WT mice (*p* = .786). There was no significant effect of stress or genotype on body weight in this sample (data not shown).

**Table 2.** Additional anatomical measures. Statistics are reported as mean ± standard deviation. *Main effect of stress, *p* < .05. †Stress × genotype interaction, *p* < .05.

|  | Wild type<br>control | Wild type<br>stress | ZnT3 KO<br>control | ZnT3 KO<br>defeated |
| --- | --- | --- | --- | --- |
| **Organ weights** | ($n = 6$) | ($n = 6$) | ($n = 6$) | ($n = 6$) |
| Spleen weight (mg)* | 74.1 ± 6.8 | 92.6 ± 16.5 | 81.1 ± 20.7 | 108.1 ± 37.6 |
| Adrenal gland<br>weight (mg)* | 4.1 ± 0.7 | 4.8 ± 0.8 | 3.2 ± 0.5 | 5.3 ± 1.3 |
| Adrenal gland<br>weight (% of body)$^\dagger$ | 0.16 ± 0.02 | 0.18 ± 0.03 | 0.13 ± 0.03 | 0.21 ± 0.04 |
| **Microglial density** | ($n = 6$) | ($n = 6$) | ($n = 6$) | ($n = 6$) |
| PFC<br>(microglia/mm$^2$) | 439.9 ± 43.7 | 444.1 ± 37.8 | 469.4 ± 45.4 | 460.0 ± 38.5 |
| dHPC<br>(microglia/mm$^2$) | 294.6 ± 41.2 | 274.6 ± 21.7 | 305.2 ± 36.8 | 290.3 ± 54.3 |
| vHPC<br>(microglia/mm$^2$) | 199.6 ± 15.2 | 195.4 ± 21.8 | 201.6 ± 20.2 | 196.0 ± 18.4 |
| BLA<br>(microglia/mm$^2$) | 435.8 ± 39.8 | 416.7 ± 13.1 | 436.0 ± 30.1 | 441.2 ± 17.0 |
| **MRI volumetric**<br>**analysis** | ($n = x$) | ($n = x$) | ($n = x$) | ($n = x$) |
| Prefrontal cortex<br>volume (mm$^3$) | 11.4 ± 0.7 | 11.7 ± 0.6 | 11.2 ± 0.8 | 11.8 ± 1.3 |
| Hippocampus<br>volume (mm$^3$) | 4.5 ± 0.3 | 4.5 ± 0.4 | 4.8 ± 0.5 | 49.4 ± 19.2 |
| Amygdala volume<br>(mm$^3$) | 9.9 ± 0.7 | 10.2 ± 0.5 | 10.2 ± 0.6 | 4.7 ± 0.5 |

### 3.3 Hippocampal cell proliferation

The number of cells in the dentate gyrus positively-labeled for the proliferation marker Ki67 was assessed in brains collected 1 day after the final episode of stress. Cell proliferation was not significantly affected by stress, nor did it differ between genotypes (Figure 5).

**Figure 5.**
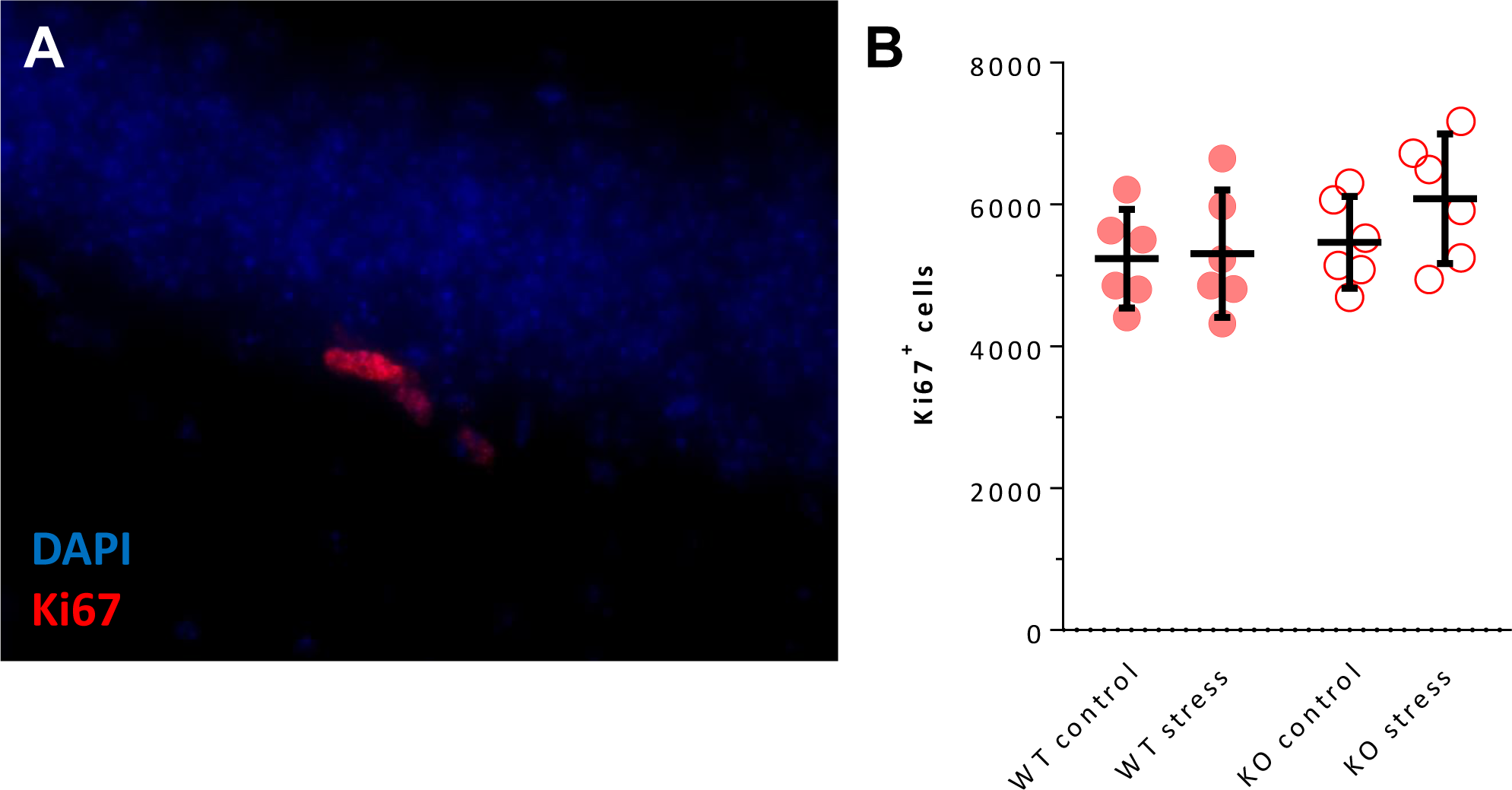
Hippocampal neurogenesis in WT and ZnT3 KO mice following repeated social defeat stress. **A.** Image of cells in the subgranular zone of the hippocampal dentate gyrus immunolabeled for the cell proliferation marker Ki67. **B.** Estimates of the total number of Ki67-positive cells in the granule cell layer and subgranular zone of the dentate gyrus, in brains collected 24 h after the last episode of defeat stress. There was no effect of stress or difference between genotypes. Error bars represent 95% CIs.

### 3.4 Microglial analysis

Using a thresholding procedure, gross changes in microglial morphology were assessed in brains collected 1 day after the final episode of stress (Figure 6). In the PFC and dHPC, there was no effect of stress or difference between genotypes. Likewise, there was no significant effect of stress or difference between genotypes in the vHPC, though there was a trend toward an interaction between stress and genotype [*F*(1,20) = 4.30, *p* = .051]. Finally, in the BLA, there was no significant effect of stress [*F*(1,20) = 4.15, *p* = .055], though the stressed mice did tend to have more Iba1^+^ labeling than the controls. There was no difference between genotypes. Next, the density of microglia was assessed (Table 2). There was no effect of stress or genotype for the PFC, dHPC, vHPC, or BLA. In summary, when microglial status was assessed, either by thresholding of Iba1 immunolabeling or by quantification of microglial density, there were no significant differences across several brain regions.

**Figure 6.**
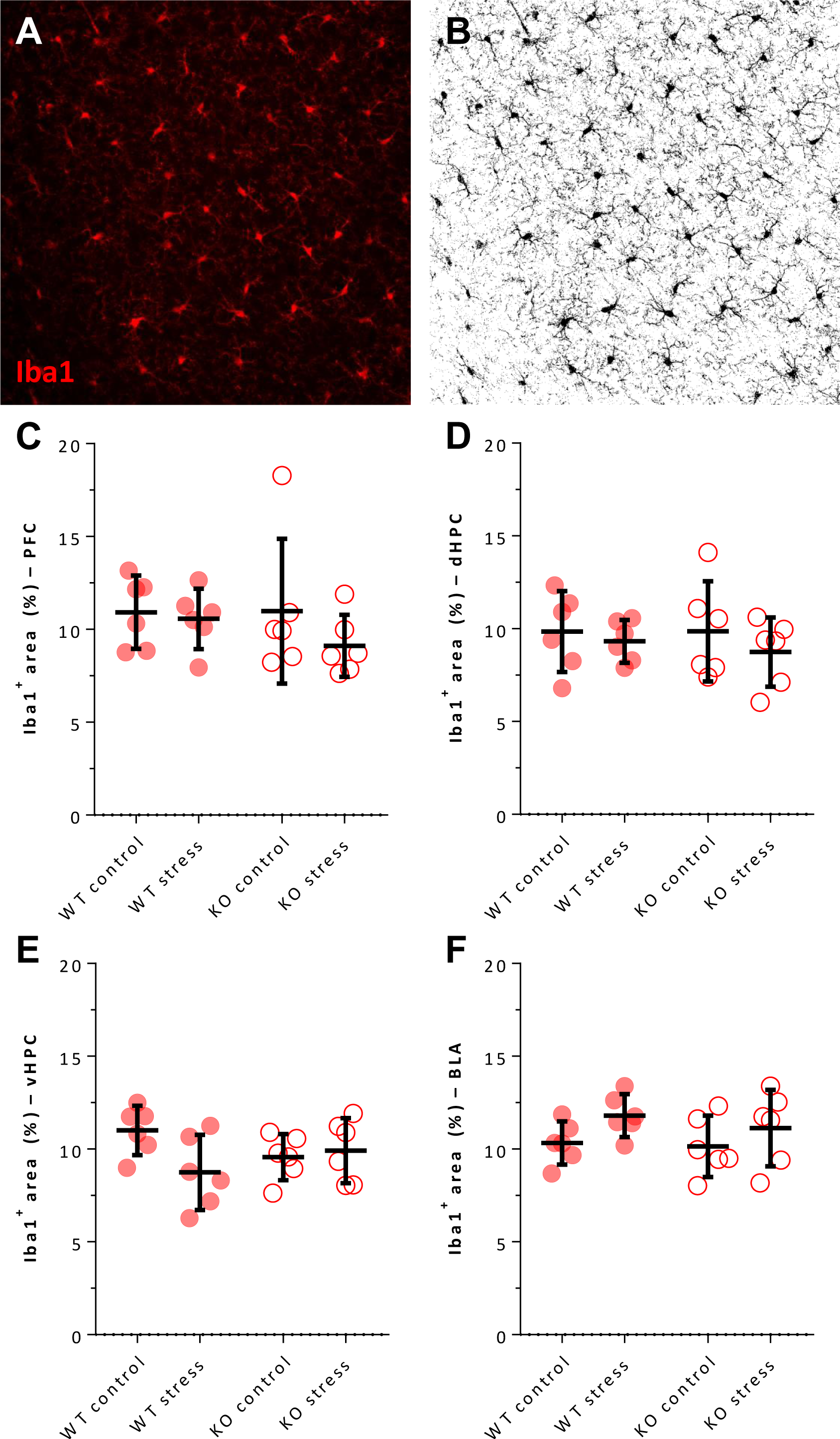
Microglial morphology in WT and ZnT3 KO mice following repeated social defeat stress. **A.** Image of microglia in the basolateral amygdala, immunolabeled for the microglial marker Iba1. **B.** The same image, after a thresholding procedure is applied to binarize the image into areas of Iba1-positive and Iba1-negative labeling. A greater percentage of positive labeling can reflect a change in microglial morphology. **C-F.** Quantification of the percentage of total area positively labeled for Iba1 across several regions of interest (prefrontal cortex, dorsal hippocampus, ventral hippocampus, and basolateral amygdala, respectively), in brains collected 24 h after the last episode of defeat stress. There was no effect of stress or difference between genotypes. Error bars represent 95% CIs.

### 3.5 MRI volumetric analysis

Volumes of several brain regions were assessed by ex-vivo MRI of brains collected 15 days after the final episode of stress (Figure 7A). First, we verified that there was no effect of stress or genotype on body weight in the subset of mice from which brains were collected (WT-control: 26.0 ± 3.1 g; WT-stress: 26.3 ± 1.9 g; KO-control: 27.2 ± 3.1 g; KO-stress: 26.9 ± 2.9 g). For the CC, the effect of stress differed based on the genotype of the mice [stress × genotype interaction: *F*(1,20) = 7.66, *p* = .009; Figure 7B], with stress decreasing CC volume by 8.2% in the ZnT3 KO mice (*p* = .002; Bonferroni-corrected) while having no effect in the WT mice (*p* = .918). It also appeared that, in the control groups, ZnT3 KO mice had larger CC volumes than WT mice. This was confirmed using a post-hoc Tukey test (WT-control vs. KO-control: *p* < .001).

**Figure 7.**
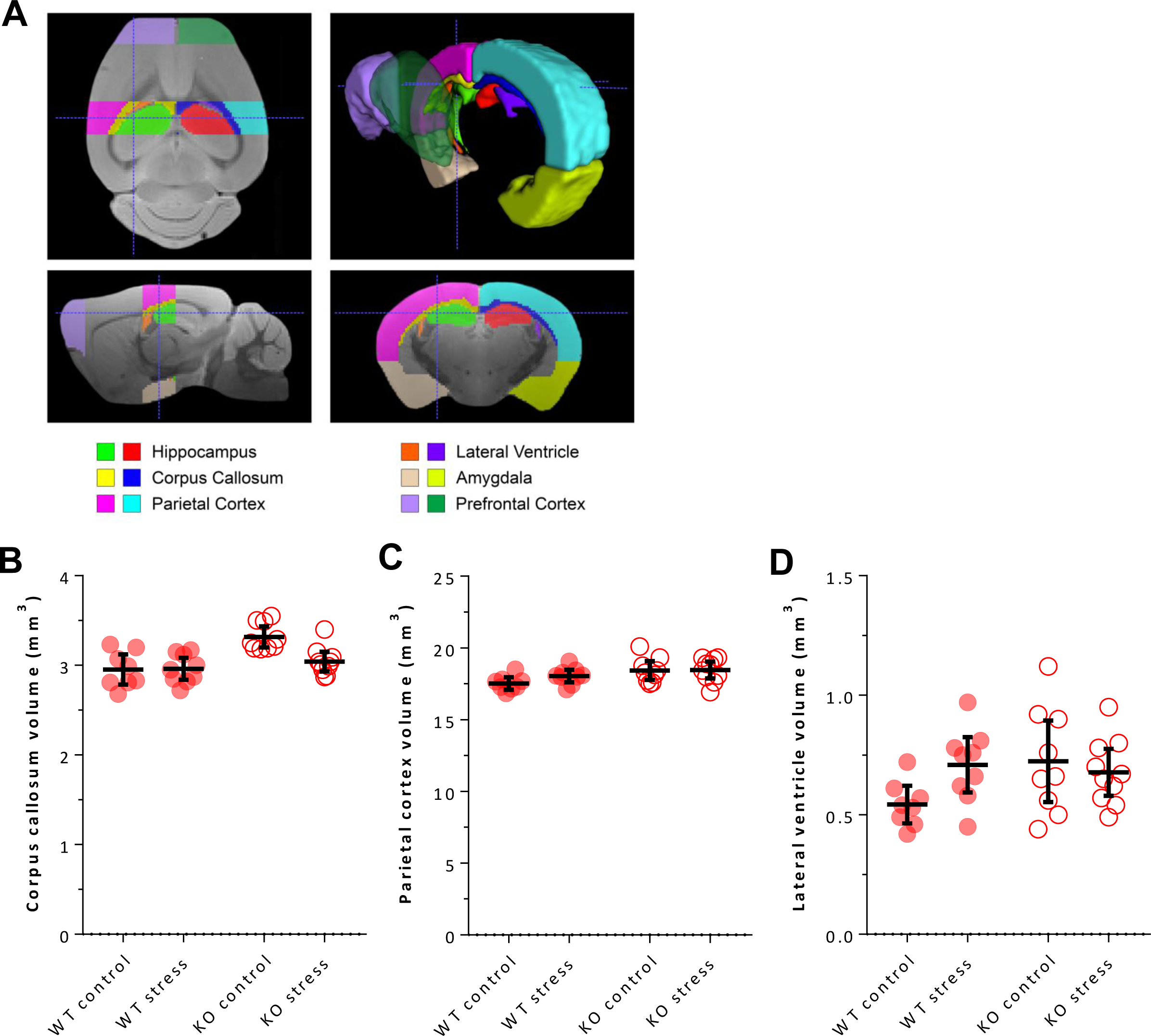
Magnetic resonance imaging volumetric analysis of several brain regions in WT and ZnT3 KO mice following repeated social defeat stress. Brains were collected 15 days after the final episode of stress. **A.** Depiction of the various regions of interest. There was no effect of stress or difference between genotypes in the volume of the prefrontal cortex, amygdala, or hippocampus **B.** Stress decreased the volume of the corpus callosum in the ZnT3 KO mice but had no effect on the WT mice. **C.** Independently of stress, ZnT3 KO mice had larger parietal cortex volumes than did WT mice. **D.** While there was no main effect of stress or genotype, the volume of the lateral ventricles was larger in the ZnT3 KO controls than in the WT controls. Error bars represent 95% CIs.

For the parietal cortex, there was no significant effect of stress, but there was a difference between genotypes [*F*(1,20) = 7.66, *p* = .009; Figure 7C], with parietal cortex being 3.6% larger in the ZnT3 KO mice than in the WT mice. For the LV, there was no significant effect of stress or genotype, though this interpretation was complicated by a trend toward an interaction between the two factors [*F*(1,20) = 3.99, *p* = .054; Figure 7D], suggesting that stress could be obscuring a difference between genotypes. We therefore compared LV volumes in the non-stressed controls. However, the difference was not significant (post-hoc Tukey test, WT-control vs. KO-control: *p* = .114). There was no effect of stress or genotype on the volume of prefrontal cortex, amygdala, or hippocampus (Table 2).

## 4. DISCUSSION

Social avoidance is a well-characterized outcome of RSD (Krishnan et al., 2007). We observed that both WT and ZnT3 KO mice became avoidant of an aggressive CD-1 mouse following RSD. Only WT mice avoided a novel conspecific, however; stressed ZnT3 KO mice did not. Given previous reports that male ZnT3 KO mice show increased (Martel et al., 2011) and decreased (Yoo et al., 2016) social interaction with a novel mouse, it is worth highlighting that we found no baseline differences in sociability; interaction time with either a conspecific or a CD-1 mouse did not differ between genotypes in non-stressed controls.

The lack of conspecific avoidance could be interpreted as a cognitive deficit, such that ZnT3 KO mice successfully learn to avoid CD-1 mice but fail to generalize the avoidance response to mice of a different strain. This interpretation fits with previous findings of mild impairments in ZnT3 KO mice, including in fear memory (Martel et al., 2010), texture discrimination (Wu & Dyck, 2018), object recognition memory (Martel et al., 2011) and spatial reversal or working memory (Cole et al., 2011; Martel et al., 2011; Sindreu et al., 2011).

On the other hand, it is arguable whether the lack of conspecific avoidance should be considered a “cognitive deficit.” From an anthropomorphic perspective, the ideal response to social stress would be to avoid the cause of the stress while *not* generalizing this response into depression-like withdrawal from all social interactions. Another interpretation of our findings, then, is that the lack of conspecific avoidance indicates resilience to stress. Defining a lack of stress-induced avoidance as a desirable trait – and, conversely, avoidance as an indicator of susceptibility to depression-like effects – is a common interpretation, for a number of reasons: 1) avoidance has obvious parallels to social withdrawal, a symptom of depression; 2) avoidance is associated with a number of other depression-like outcomes (Krishnan et al., 2007), including reduced sucrose preference, altered circadian function, and decreased body weight; 3) avoidance can be reversed by chronic treatment with antidepressant drugs (Tsankova et al., 2006).

Because resilience is usually defined by interaction with a CD-1 mouse – rather than a conspecific – it is somewhat difficult to directly compare the “resilience” observed by ZnT3 KO mice in our study to the “resilience” reported in many other studies. One way to further address this issue would be to examine whether ZnT3 KO mice are resilient to other depression-like behaviours. Sucrose preference or circadian function would be good candidates; unfortunately, neither was assessed in the current study. We examined anxiety, but this does little to clarify the matter, because increased anxiety can occur independently from social avoidance (Krishnan et al., 2007); that is, mice can become “anxious” without being susceptible to depression-like behaviours. We did assess body weight, which provides information about susceptibility to depression-like effects, as both weight gain and weight loss are symptoms of depression. However, we did not observe an effect of stress. This is perhaps not surprising, as the effect of RSD on body weight is quite variable between studies, with reports of weight loss or attenuated weight gain (Kudryavtseva et al., 1991; Krishnan et al., 2007; Venzala et al., 2012) as well as increased weight gain (Bartolomucci et al., 2004; Dubreucq et al., 2012).

We also examined how stress affects cognition in ZnT3 KO mice. RSD had no significant effect on contextual fear memory, but the results of the cued fear memory test were more interesting. Stress enhanced cued fear memory in ZnT3 KO mice, but WT mice were not affected. Interpreting this effect is complicated, however, because the two genotypes did not start out from the same baseline; under control conditions, ZnT3 KO mice showed weaker fear memory than WT mice. We also observed ZnT3 KO mice to have weaker memory in the contextual fear test, supporting the results of Martel et al. (2010), who previously showed fear memory impairments in these mice.

The lack of a strong effect of stress on fear memory was somewhat surprising, considering that others have observed enhanced fear memory after chronic restraint in rats (Conrad et al., 1999; Suvrathan et al., 2014) and RSD in mice (Fuertig et al., 2016; Lisboa et al., 2018). It is possible that the effect did not persist over the 10-day gap between RSD and fear memory testing in the present study. We also failed to detect an effect of RSD on memory in a spatial Y-maze test, despite the well-documented deleterious effect of chronic stress on spatial memory (Conrad et al., 1996; Conrad, 2010; Wang et al., 2011). It might be the case that a longer period of stress is required; at 10 days, the duration of stress in our experiment was relatively short. And the period of time between RSD and testing is, again, a possible factor. The hippocampal atrophy that occurs in response to 21 days of restraint stress recovers within 5-10 days (Conrad et al., 1999). It is possible that any memory deficits produced by stress in our experiment recovered over the 8 days between RSD and the Y-maze test.

Chronic social stress decreases the proliferation, survival, differentiation, and maturation of adult-born cells in the hippocampus, at least at certain time points (Czéh et al., 2001; Van Bokhoven et al., 2011; Chen et al., 2015; McKim et al., 2016). Further, some behavioural effects of RSD are mediated by changes in neurogenesis (Lagace et al., 2010; Lehmann et al., 2013). ZnT3 KO mice fail to show the increase in neurogenesis that is normally seen following hypoglycemia (Suh et al., 2009), and we have observed that these mice also do not show the increase in neurogenesis that normally results from enriched housing (Chrusch et al., unpublished). We therefore speculated that the effect of stress on neurogenesis might also be abnormal in ZnT3 KO mice, and that this might account for the altered behavioural profile. We examined cell proliferation 24 h after RSD, but found it to be unaffected by stress, regardless of genotype. Though unexpected, this is consistent with previous findings that the number of proliferating cells in S-phase is decreased immediately after the completion of 10 days of RSD, but not 24 h later (Lagace et al., 2010).

We also examined the status of microglia. RSD “activates” microglia, and this is associated with the development of anxiety-like behaviour (Wohleb et al., 2011, 2014; McKim et al., 2018). While we are unaware of direct evidence that microglia function abnormally in ZnT3 KO mice, microglia do express receptors that are sensitive to modulation by zinc, such as P2X7 receptors (Liu et al., 2008). Further, pretreating cultured microglia with zinc prior to stimulation by lipopolysaccharide increases the release of pro-inflammatory cytokines (Higashi et al., 2017). We assessed microglial morphology using a thresholding procedure that has previously been effective at detecting RSD-induced changes (Wohleb et al., 2011, 2014; McKim et al., 2018). However, we were unable to detect an effect of stress on microglial morphology or density, despite the development of anxiety-like behaviour. The RSD protocol that has been found to induce microglial activation involves introducing a dominant CD-1 intruder into a cage of three mice for 2 h. It is possible that this is more stressful than the protocol used in the current study, and likely results in greater wounding and inflammation, which may contribute to microglial activation.

Finally, we used MRI to conduct a volumetric analysis. CC volume was greater in control ZnT3 KO mice than in WT mice, and stress decreased CC volume only in ZnT3 KO mice. To our knowledge, this is the first indication of white matter abnormalities in ZnT3 KO mice. Interestingly, it has previously been observed that the CC is larger in mice that are resilient to RSD than in susceptible mice (Anacker et al., 2016). Assuming that a larger CC is a protective factor, this could explain why ZnT3 KO mice show diminished social avoidance, though it does not explain why CC volume was reduced by stress only in ZnT3 KO mice. We also found that parietal cortex was larger in ZnT3 KO mice than in WT mice, which supports a finding of increased cortical size by Yoo et al. (2017). One limitation is that our analysis included portions, but not the entirety, of several structures – most notably, our definition of the hippocampus was limited to the dorsal region. And there are inherent challenges to accurately delineating regions of interest on MRI, as was particularly the case for the amygdala in the present study. In future studies, it would be valuable to apply complementary methods of quantifying regional volumes, as well as other MRI methods, such as diffusion MRI, for assessing inter- and intra-structural connectivity.

## 5 CONCLUSIONS

Our primary aim was to examine how chronic stress, in the form of RSD, impacts mice that lack vesicular zinc due to genetic deletion of ZnT3. We found that these mice, unlike WT mice, did not become avoidant of a novel conspecific, suggesting increased resilience to the depression-like effects of stress. ZnT3 KO mice were not entirely unaffected by stress, however; they did become avoidant of a CD-1 mouse and also exhibited stress-induced anxiety-like behaviour. Finally, cued fear memory was enhanced by stress in ZnT3 KO mice, but not in their WT counterparts. Thus, a lack of vesicular zinc modulates the outcomes of RSD, but not in a straightforward fashion. We were unable to account for these behavioural effects through differences in hippocampal neurogenesis or microglial activation. However, we did observe ZnT3 KO mice to have larger CC volumes than WT mice. Further study will be required to determine whether this neuroanatomical abnormality is protective against the depression-like effects of RSD.

## 6. ACKNOWLEDGEMENTS

The authors thank Abril Valverde Rascón, Katy Sandoval, and Angela Pochakom for their assistance. The authors also acknowledge the facilities provided by the HBI Advanced Microscopy Platform (Calgary, AB, Canada) and Dr. Vincent Ebacher for his assistance. Finally, the authors acknowledge the facilities, and the scientific and technical assistance, of the National Imaging Facility at the Florey Institute of Neuroscience and Mental Health (Parkville, VIC, Australia).

## Supplemental Methods

### Immunofluorescence labeling

For immunofluorescence labeling, the procedure was as follows: 3 × 10 min wash in PBS; blocking for 1 h in PBS containing 0.3% Triton X-100 (PBSx) and 4% normal goat serum (NGS); incubation overnight at room temperature in PBSx containing 2% NGS and the primary antibody (1:1000 rabbit anti-Iba1, Wako #019-19741; or 1:2000 rabbit anti-Ki67, Leica NCL-Ki67p); 3 × 10 min wash in PBSx; incubation overnight at room temperature in PBSx containing the secondary antibody (1:1000, biotin-conjugated goat anti-rabbit, Jackson ImmunoResearch 111-065-144); 3 × 10 min wash in PBSx; incubation for 1 h in PBSx containing the tertiary antibody (1:1000, Alexa-Fluor 594-conjugated streptavidin, Jackson ImmunoResearch 016-580-084) with 4′,6-diamidino-2-phenylindole (DAPI; 1:1000) added for the final 15 min; 3 × 10 min wash in PBS.

**Table S1.** ANOVA results.

|  | D.F. | Stress | Genotype | Stress × Genotype |
| --- | --- | --- | --- | --- |
| **Social interaction test** |  |  |  |  |
| Interaction time – empty cage | 1, 82 | **$F = 4.92, p = .029$** | $F = 0.58, p = .450$ | $F = 0.13, p = .719$ |
| Interaction time – conspecific | 1, 82 | **$F = 23.20, p < .001$** | $F = 1.39, p = .243$ | **$F = 6.53, p = .012$** |
| Interaction time – CD-1 | 1, 82 | **$F = 22.53, p < .001$** | $F = 0.04, p = .835$ | $F = 0.26, p = .608$ |
| Corner time – empty cage | 1, 82 | $F = 1.66, p = .201$ | $F = 0.60, p = .440$ | $F = 0.63, p = .430$ |
| Corner time – conspecific | 1, 82 | **$F = 25.74, p < .001$** | $F = 1.81, p = .182$ | **$F = 6.36, p = .014$** |
| Corner time – CD-1 | 1, 82 | **$F = 37.18, p < .001$** | **$F = 4.12, p = .046$** | $F = 2.27, p = .136$ |
| Interaction ratio – CD-1 | 1, 81 | **$F = 8.32, p = .005$** | $F = 0.09, p = .762$ | $F < 0.01, p = .948$ |
| Distance traveled – empty cage | 1, 82 | $F = 0.33, p = .566$ | $F < .01, p = .953$ | $F = 0.24, p = .624$ |
| Immobility time – conspecific | 1, 82 | **$F = 38.72, p < .001$** | $F = 0.05, p = .826$ | $F = 2.45, p = .121$ |
| Immobility time – CD-1 | 1, 82 | **$F = 19.53, p < .001$** | $F = 1.54, p = .219$ | $F = 0.04, p = .840$ |
| **Elevated plus-maze** |  |  |  |  |
| Open arm time | 1, 58 | **$F = 8.74, p = .004$** | $F = 2.46, p = .122$ | $F = 0.54, p = .465$ |
| Center time | 1, 58 | **$F = 10.99, p = .002$** | $F = 3.21, p = .078$ | $F = 0.77, p = .384$ |
| Distance traveled | 1, 58 | $F = 0.97, p = .328$ | $F < 0.01, p = .983$ | $F = 2.23, p = .141$ |
| **Novelty-suppressed feeding** |  |  |  |  |
| Latency to feed (s) – novel cage | 1, 58 | **$F = 40.43, p < .001$** | $F = 0.27, p = .606$ | $F = 0.09, p = .764$ |
| Latency to feed (s) – home cage | 1, 58 | $F = 0.56, p = .459$ | $F = 0.54, p = .467$ | $F = 0.01, p = .918$ |
| Food consumption (g) | 1, 58 | $F = 1.06, p = .307$ | $F = 3.68, p = .060$ | $F = 0.66, p = .419$ |
| Weight loss (g) | 1, 58 | **$F = 42.65, p < .001$** | $F = 2.47, p = .121$ | $F = 0.05, p = .828$ |
| **Y-maze spatial memory** |  |  |  |  |
| Novel arm entries (% of total entries) | 1, 58 | $F = 0.56, p = .458$ | $F = 0.35, p = .554$ | $F = 0.27, p = .603$ |
| Novel arm time (s) | 1, 58 | $F = 0.89, p = .348$ | $F = 0.01, p = .943$ | $F = 0.05, p = .830$ |
| Total arm entries | 1, 58 | $F = 0.12, p = .726$ | $F = 0.16, p = .689$ | **$F = 4.18, p = .045$** |
| **Conditioned fear** |  |  |  |  |
| Freezing time – training pre-cue | 1, 58 | $F < 0.01, p = .969$ | $F = 0.10, p = .760$ | $F = 0.41, p = .523$ |
| Freezing time – training during cue | 1, 58 | $F = 1.77, p = .189$ | $F = 0.70, p = .406$ | $F = 0.23, p = .632$ |
| Freezing time – training post-cue | 1, 58 | $F = 0.14, p = .705$ | $F = 0.05, p = .825$ | $F = 0.01, p = .918$ |
| Freezing time – pre-cue | 1, 58 | $F = 0.82, p = .369$ | $F = 0.05, p = .824$ | $F = 0.32, p = .577$ |
| Freezing time – cued fear | 1, 58 | $F = 3.88, p = .054$ | $F = 0.99, p = .324$ | **$F = 5.89, p = .018$** |
| Freezing time – contextual fear | 1, 58 | $F = 3.02, p = .087$ | **$F = 6.65, p = .012$** | $F = 0.55, p = .463$ |
| **Body and organ weights** |  |  |  |  |
| Weight change – pre- to post-RSD | 1, 77 | $F = 0.24, p = .627$ | $F = 0.04, p = .835$ | $F = 0.01, p = .933$ |
| Spleen weight | 1, 20 | **$F = 5.76, p = .026$** | $F = 1.40, p = .250$ | $F = 0.20, p = .660$ |
| Spleen weight<br>(% of body weight) | 1, 20 | **$F = 5.40, p = .031$** | $F = 2.71, p = .116$ | $F = 0.76, p = .394$ |
| Adrenal weight | 1, 20 | **$F = 15.86, p < .001$** | $F = 0.25, p = .662$ | $F = 3.92, p = .062$ |
| Adrenal weight<br>(% of body weight) | 1, 20 | **$F = 16.32, p < .001$** | $F < 0.01, p = .985$ | **$F = 7.87, p = .011$** |
| Body weight | 1, 20 | $F = 1.00, p = .328$ | $F = 1.16, p = .294$ | $F = 0.56, p = .463$ |
| **Hippocampal cell proliferation** |  |  |  |  |
| Ki67<sup>+</sup> cells | 1, 20 | $F = 1.23, p = .281$ | $F = 2.62, p = .121$ | $F = 0.77, p = .391$ |
| **Microglial activation** |  |  |  |  |
| Iba1<sup>+</sup> area – PFC | 1, 20 | $F = 1.34, p = .261$ | $F = 0.53, p = .476$ | $F = 0.62, p = .440$ |
| Iba1<sup>+</sup> area – dorsal HPC | 1, 20 | $F = 1.05, p = .319$ | $F = 0.12, p = .728$ | $F = 0.14, p = .718$ |
| Iba1<sup>+</sup> area – ventral HPC | 1, 20 | $F = 2.32, p = .143$ | $F = 0.05, p = .831$ | $F = 4.30, p = .051$ |
| Iba1<sup>+</sup> area – BLA | 1, 20 | $F = 4.15, p = .055$ | $F = 0.49, p = .490$ | $F = 0.16, p = .692$ |
| Iba1<sup>+</sup> cell density – PFC | 1, 20 | $F = 0.02, p = .881$ | $F = 1.79, p = .196$ | $F = 0.16, p = .690$ |
| Iba1<sup>+</sup> cell density – dorsal HPC | 1, 20 | $F = 1.13, p = .300$ | $F = 0.64, p = .433$ | $F = 0.03, p = .877$ |
| Iba1<sup>+</sup> cell density – ventral HPC | 1, 20 | $F = 0.39, p = .537$ | $F = 0.03, p = .872$ | $F = 0.01, p = .926$ |
| Iba1<sup>+</sup> cell density – BLA | 1, 20 | $F = 0.39, p = .539$ | $F = 1.24, p = .279$ | $F = 1.22, p = .283$ |
| **MRI volumetric analysis** |  |  |  |  |
| PFC volume | 1, 32 | $F = 1.88, p = .180$ | $F = 0.02, p = .897$ | $F = 0.34, p = .563$ |
| Hippocampus volume | 1, 32 | $F = 0.28, p = .603$ | $F = 2.39, p = .132$ | $F = 0.57, p = .457$ |
| Amygdala volume | 1, 32 | $F = 1.39, p = .251$ | $F = 1.25, p = .271$ | $F = 0.01, p = .943$ |
| Lateral ventricle volume | 1, 32 | $F = 1.31, p = .261$ | $F = 1.90, p = .177$ | $F = 3.99, p = .054$ |
| Corpus callosum volume | 1, 32 | **$F = 5.66, p = .024$** | **$F = 16.07, p < .001$** | **$F = 6.40, p = .017$** |
| Parietal cortex volume | 1, 32 | $F = 1.37, p = .251$ | **$F = 7.66, p = .009$** | $F = 1.06, p = .311$ |
| Body weight | 1, 32 | $F < 0.01, p = .986$ | $F = 0.94, p = .339$ | $F = 0.09, p = .768$ |

## Abbreviations

BLA: basolateral amygdala
CC: corpus callosum
dHPC: dorsal hippocampus
EPM: elevated plus-maze
LV: lateral ventricles
NAc: nucleus accumbens
NSF: novelty-suppressed feeding
PBS: phosphate-buffered saline
PFA: paraformaldehyde
PFC: prefrontal cortex
RSD: repeated social defeat
vHPC: ventral hippocampus
ZnT3: zinc transporter 3

